# ASTRO: Automated Spatial Whole-Transcriptome RNA-Expression Workflow

**DOI:** 10.1101/2025.01.24.634814

**Authors:** Dingyao Zhang, Zhiyuan Chu, Yiran Huo, Zhiliang Bai, Rong Fan, Jun Lu, Mark Gerstein

## Abstract

**Motivation:** Despite significant advances in spatial transcriptomics, the analysis of formalin-fixed paraffin-embedded (FFPE) tissues, which constitute most clinically available samples, remains challenging. Additionally, capturing both coding and noncoding RNAs in a spatial context poses significant challenges. We recently introduced Patho-DBiT, a technology designed to address these unmet needs. However, the marked differences between Patho-DBiT and existing spatial transcriptomics protocols necessitate specialized computational tools for comprehensive whole-transcriptome analysis in FFPE samples.

**Results:** Here, we present ASTRO, an automated pipeline developed to process spatial transcriptomics data. In addition to supporting standard datasets, ASTRO is optimized for whole-transcriptome analyses of FFPE samples, enabling the detection of various RNA species, including non-coding RNAs such as miRNAs. To compensate for the reduced RNA quality in FFPE tissues, ASTRO incorporates a specialized filtering step and optimizes spatial barcode calling, increasing the mapping rate. These optimizations allow ASTRO to spatially quantify coding and non-coding RNA species in the entire transcriptome and achieve robust performance in FFPE samples.

**Availability:** Codes are available at GitHub (https://github.com/gersteinlab/ASTRO).

## 1 Introduction

With spatial information incorporated, spatial transcriptomics technologies have revolutionized transcriptomic analyses in recent years, opening a new era of genomics research (Bressan *et al*., 2023; Deng *et al*., 2023; Chen *et al*., 2024; Baysoy *et al*., 2023). Although the field has made remarkable strides, most spatial transcriptomics methods still focus on mRNAs and do not capture the entire transcriptome. Yet, extensive evidence shows that various noncoding RNAs play critical biological roles in tissues, underscoring the importance of spatially profiling these molecules (Chen and Kim, 2024; Mattick *et al*., 2023). Furthermore, formalin-fixed paraffin-embedded (FFPE) tissues are commonly used in hospital pathology departments, and the extensive collections of FFPE blocks represent an invaluable resource for translational research (Blow, 2007). However, sequencing these samples faces significant challenges, including RNA fragmentation, degradation, chemical modifications, and the loss of poly-A tails, especially when stored under suboptimal conditions(Levin *et al*., 2020).

To address this gap, we recently developed **patho**logy-compatible **d**eterministic **b**arcoding **i**n **t**issue (Patho-DBiT) (Bai *et al*., 2024), which leverages *in situ* polyadenylation to enable spatial whole-transcriptome sequencing in clinically archived FFPE tissues. However, the significant differences between traditional mRNA sequencing and whole-transcriptome sequencing, as well as between fresh-frozen samples and FFPE samples, necessitate a specialized computational pipeline to facilitate comprehensive spatial profiling of whole transcriptomics in these clinical-level tissues.

In this study, we develop and implement ASTRO: (**A**utomated **S**patial Whole-**T**ranscriptome **R**NA-Expression w**O**rkflow), a spatial transcriptomics mapping pipeline optimized for both coding and noncoding RNAs (ncRNAs) as well as FFPE samples (Fig. 1). The pipeline employs a scoring system to capture ncRNAs at different maturation stages and removes incorrect or non-expressed ncRNA annotations, enabling robust spatial profiling of the full spectrum of ncRNAs. Additionally, ASTRO distinguishes intron reads from exon reads, adding further depth to RNA biology analyses. Because FFPE samples often exhibit severe RNA degradation(Levin *et al*., 2020), ASTRO maximize the information obtained from each sample by tolerating insertions and deletions (indels) and variations in barcode regions during demultiplexing. Subsequently, ASTRO deploys a post-alignment filter to eliminate invalid reads. By integrating these advanced features, we developed a robust pipeline specifically tailored for spatial whole transcriptome analysis in FFPE tissues.

**Fig 1.**
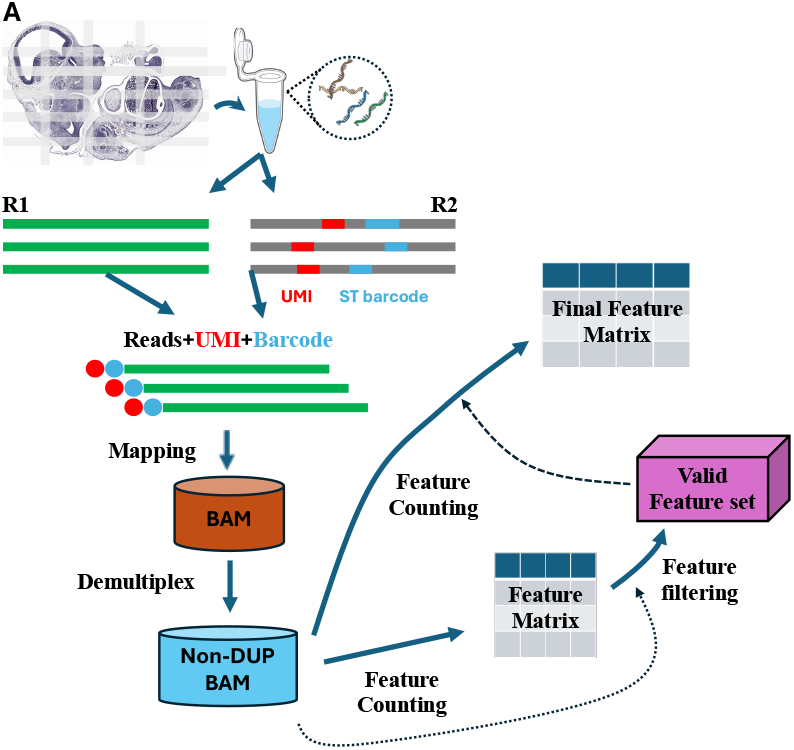
Workflow of ASTRO. **(A)** A schematic overview of the ASTRO workflow is shown. Spatial transcriptomics data typically comprise paired FASTQ files: R1, containing transcriptome RNA sequences, and R2, containing spatial barcodes and UMIs. The pipeline merges R2 information with R1 to create a combined FASTQ file, which is then mapped to the genome. After mapping, demultiplexing is performed on the resulting BAM file to generate a non-duplicate (Non-DUP) BAM file for feature counting. An intermediate feature matrix is then generated and analyzed alongside the non-duplicate BAM file to produce a valid feature set. Finally, the valid feature set is used in a second round of feature counting to generate the final feature matrix.

## 2 Methods

### 2.1 The ASTRO Pipeline

An overview of the ASTRO pipeline is presented in **Figure 1**. ASTRO takes FASTQ files as input and produces a gene-pixel (feature-location) matrix. The pipeline supports various RNA species—including mRNAs, lncRNAs, tRNAs, and miRNAs. Because of the biological differences between introns and exons, ASTRO separates intron reads and exon reads by default, facilitating downstream analyses such as RNA velocity. In addition to generating the final matrix, ASTRO outputs intermediate files (e.g., BAM files) for further analyses. The ASTRO pipeline is available as both a bash pipeline and a Python package in the Gerstein Lab GitHub repository: https://github.com/gersteinlab/ASTRO.

### 2.2 Demultiplexing and genomic alignment in ASTRO

During demultiplexing, the pipeline first utilizes the structure of read 2 (R2) to guide processing. It then applies Cutadapt to extract sequences corresponding to unique molecular identifiers (UMIs) and potential spatial barcodes(Martin, 2011). Because R2 may contain indels, the pipeline extracts sequences from an expanded region for spatial barcodes. To handle potential sequencing errors, ASTRO leverages Bowtie2 or STAR to build a reference of valid spatial barcodes and align the spliced R2 spatial barcode sequences to this reference, thereby determining each read’s spatial barcode(Dobin *et al*., 2013; Langmead and Salzberg, 2012). All spatial barcodes were pre-designed and therefore defined in sequence. During barcode mapping, only the best match is retained, and reads mapping equally well to multiple barcodes are discarded as ambiguous reads.

### 2.3 Feature counting in ASTRO

#### 2.3.1 Establishment of a non-coding RNA reference

For genome mapping in various research projects, the GENCODE database is among the most widely used resources(Mudge *et al*., 2025). However, its non-coding RNA annotation remains incomplete. It omits certain non-coding RNA species and, in other cases, lacks sufficient detail (e.g., mature 5p/3p miRNA isoforms and codon differences among tRNAs). To address these gaps, we created specialized GTF files for our pipeline by compiling genomic data from multiple databases, including GENCODE, miRbase, piRNAdb, GtRNAdb, and RNAcentral(Chan and Lowe, 2016; Wang *et al*., 2019; Kozomara and Griffiths-Jones, 2011; Sweeney *et al*., 2021; Mudge *et al*., 2025). We then merged similar records into single entries to form comprehensive GTF files. Further details on this procedure are available on GitHub. We produced these specialized GTF files for two genome versions: mouse mm39 and human GRCh38.

#### 2.3.2 Assign reads with overlapping annotations

For RNA fragments mapping to multiple overlapping annotations, an ‘overlap score’ was calculated using the formula (*L*_*o*_−*L*_*no*_)/*L*_*a*_, where *L*_*o*_ denotes the overlapped length between the query RNA fragment and an annotation, *L*_*no*_ represents the non-overlapped length of the query RNA fragment, and *L*_*a*_ is the total length of the genomic annotation. The annotation with the highest overlap score was considered the true annotation for the RNA fragment. In our specialized GTF, exon and transcripts are marked as different records, allowing the method to differentiate between exon and intron features. If a read has the highest overlap score with an exon feature, it is classified as exon mapping. Conversely, if a read has the highest overlap score with an intron feature, it is classified as mapped to an intron.

#### 2.3.3 Adjustment of valid features

The previous assignment step depends heavily on high-quality GTF files, as invalid entries can lead to an excessive number of spurious features in the gene expression matrix. However, due to the strong tissue specificity of many non-coding RNAs(Statello *et al*., 2020; Ludwig *et al*., 2016), it is necessary to adjust the GTF files for each dataset. The main principle behind this adjustment is that a genuine RNA structure typically displays a significantly higher read depth than its background regions, whereas a structure formed by randomly fragmented background RNA should not exhibit a substantial change in read depth. To implement this principle, we conduct an examination, including a statistical test, for each feature. Further details on this validation process are provided in the **Supplementary Materials**.

### 2.4 Evaluating the performance of ASTRO

#### 2.4.1 Collection of spatial transcriptome datasets

To assess the performance of ASTRO, we used four publicly available spatial transcriptome datasets from our previous study(Bai *et al*., 2024). The first dataset is a clinical extranodal marginal zone lymphoma of mucosa-associated lymphoid tissue (MALT) tumor biopsy, featuring 10,000 spots (or pixels) at a 20 µm pixel size. The second dataset is an FFPE biopsy of a healthy donor lymph node. The other two datasets are replicates of embryonic day 13 mouse embryo FFPE sections at a 50 µm pixel size; these two replicates were collected from two adjacent slides.

#### 2.4.2 Comparison between ASTRO and existing methods

To evaluate the impact of ASTRO on downstream analyses, we compared it against existing spatial transcriptomics pipelines. Currently, two widely used pipelines are the ST-pipeline and the Space Ranger(Navarro *et al*., 2017; Zheng *et al*., 2017); however, Space Ranger is only compatible with 10x Genomics data. Consequently, we restricted our benchmarking to the ST-pipeline (version 1.8.1) which we installed via pip. We evaluated pipeline performance on downstream analyses using two approaches. First, we aligned the clustering results with hematoxylin and eosin (H&E) staining to compare tissue structures; a more refined pipeline should capture more detailed structures. Second, we employed three quantitative metrics previously used in single-cell clustering evaluations—the Silhouette score, the Calinski–Harabasz index, and the Davies–Bouldin index— to assess clustering performance (Yu *et al*., 2022; Møller and Madsen, 2023; Jiang *et al*., 2018; Leng *et al*., 2022). Note that higher Silhouette and Calinski–Harabasz scores indicate superior clustering performance, while lower Davies–Bouldin index scores indicate better performance. This assessment was conducted on both the full dataset and a 50% sub-sampled dataset. Further details on the downsampling process are provided in the **Supplementary Materials**.

## 3 Results

To demonstrate the performance of ASTRO in whole transcriptome, we deployed it across all four spatial transcriptome samples. ASTRO successfully captured a wide range of RNA species across the datasets, including lncRNAs, miRNAs, protein-coding RNAs (mRNAs), rRNAs, scaRNAs, snoRNAs, snRNAs, tRNAs, vault RNAs, Y RNAs, and miscRNAs (**Figure 2A–2D**). Because piRNAs are highly germline-tissue-specific, we excluded them from these statistics(Wang *et al*., 2024). Beyond its broad RNA coverage, ASTRO filters out features that are not truly expressed. For instance, in the MALT sample, ASTRO identified various miRNAs while dismissing those inflated by background noise. Reads mapped to hsa-miR-4454 and hsa-miR-1260b were significantly enriched above background, indicating their validity (**Figure 2E**). In contrast, reads mapped to hsa-mir-3648 and hsa-mir-4449 were not significantly enriched, suggesting that they should not be assigned to these miRNAs (**Figure 2F**). Moreover, ASTRO captures isoform-level differences. Many miRNAs have two mature isoforms (5p isoform and 3p isoform), and the expression patterns of these isoforms are known to be different. In the MALT dataset, hsa-miR-146a-5p, hsa-miR-29b-3p, and hsa-miR-150a-5p were identified as valid features in this dataset, whereas their corresponding isoforms (hsa-miR-146a-3p, hsa-miR-29b-5p, and hsa-miR-150a-3p) should be excluded in this sample **(Figure 2G)**.

**Fig 2.**
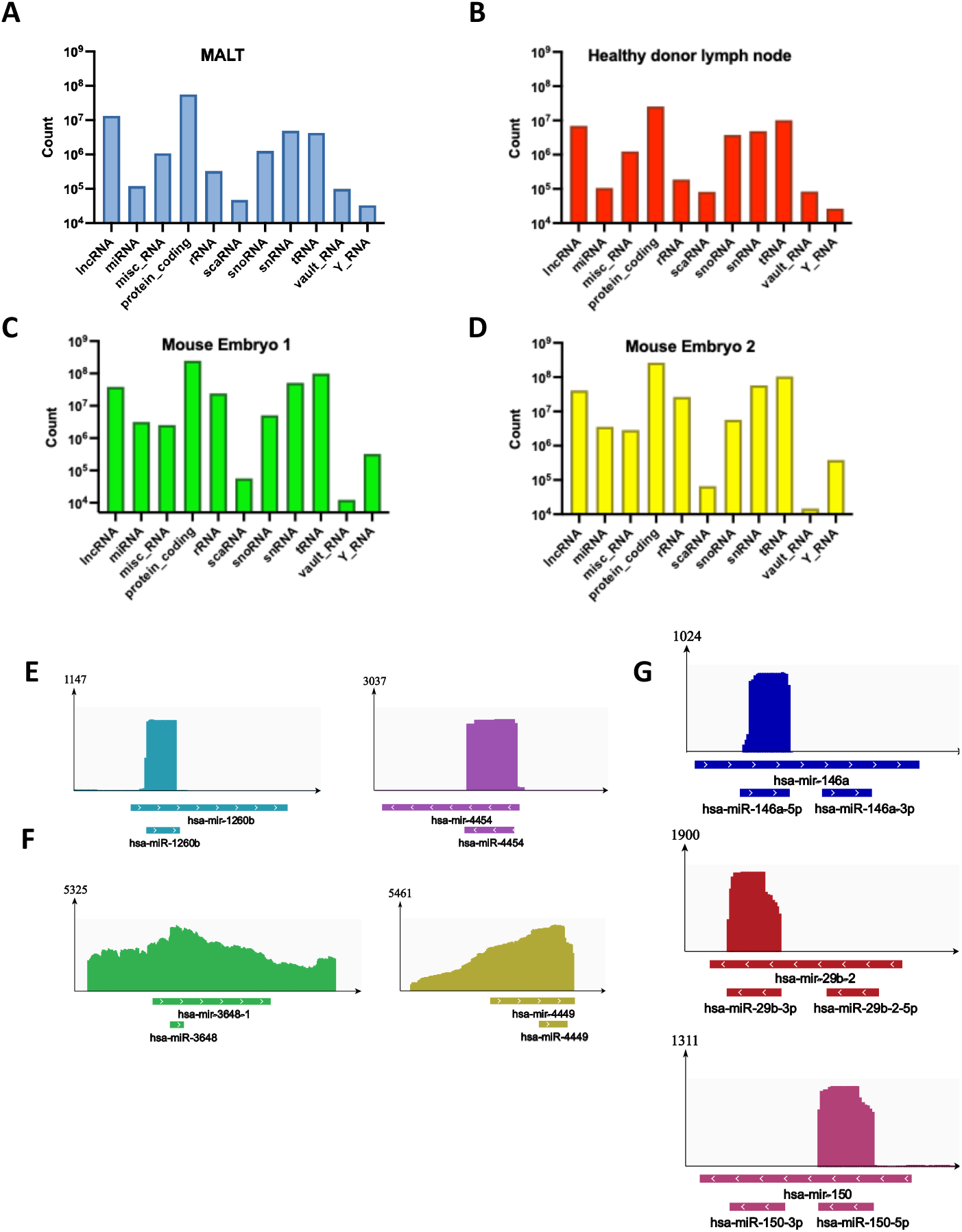
ASTRO enables whole transcriptome analysis, including miRNAs. **(A-D)** Bar plots show the ability of ASTRO to detect various RNA species, with the y-axis indicating the number of reads assigned to each species. (MALT, marginal zone lymphoma of mucosa-associated lymphoid tissue). **(E)** Examples of miRNAs, including hsa-miR-4454 and hsa-miR-1260b, that are significantly enriched compared to background levels. **(F)** Examples of miRNAs, including hsa-miR-3648 and hsa-miR-4449, that are not significantly enriched above background levels. **(G)** Examples of miRNAs, including hsa-miR-146a-5p, hsa-miR-29b-3p, and hsa-miR-150a-5p, whose isoforms display distinct patterns.

When we applied both ASTRO and the ST-pipeline to the same MALT dataset, the total number of captured reads differed. A key feature of ASTRO is its ability to distinguish between exons and introns, allowing it to capture more reads overall. In both exon-assigned and intron-assigned counts, ASTRO detected a higher gene/UMI count than ST-pipeline which does not distinguish introns from exons. To further compare the performance of ASTRO and the ST-pipeline, we applied each pipeline separately to the MALT sample. For the Silhouette score, Calinski–Harabasz index, and Davies–Bouldin index, ASTRO-based analysis achieved values of 0.1525, 1,726.9187, and 1.4279, respectively, outperforming ST-pipeline-based analysis, which produced values of 0.0989, 985.0426 and 1.5817.

Overall, ASTRO exhibited a much lower noise level, as indicated by spatial clustering and statistical tests (**Figure 3B**). Additionally, ASTRO identified detailed tissue structures that ST-pipeline overlooked (**Figures 3C, 3D**). For example, in the purple-circled region, the H&E image revealed a distinct vasculature stripe surrounded by tumor B cells. ASTRO clearly delineated this specific cell group, whereas ST-pipeline improperly split the region into multiple clusters. Similarly, in the red-circled region, ASTRO accurately distinguished a smooth muscle cell pattern from the neighboring cells, whereas ST-pipeline failed to identify this structure. Finally, we assessed all four samples using the Silhouette score, Calinski– Harabasz index, and Davies–Bouldin index. For all datasets, ASTRO-based results demonstrated superior clustering performance (**Supplementary Materials**). When subsampling was performed, ASTRO outperformed ST-pipeline across nearly all metrics and samples. (**Figure 3E, Figure S1A, S1B**).

**Fig 3.**
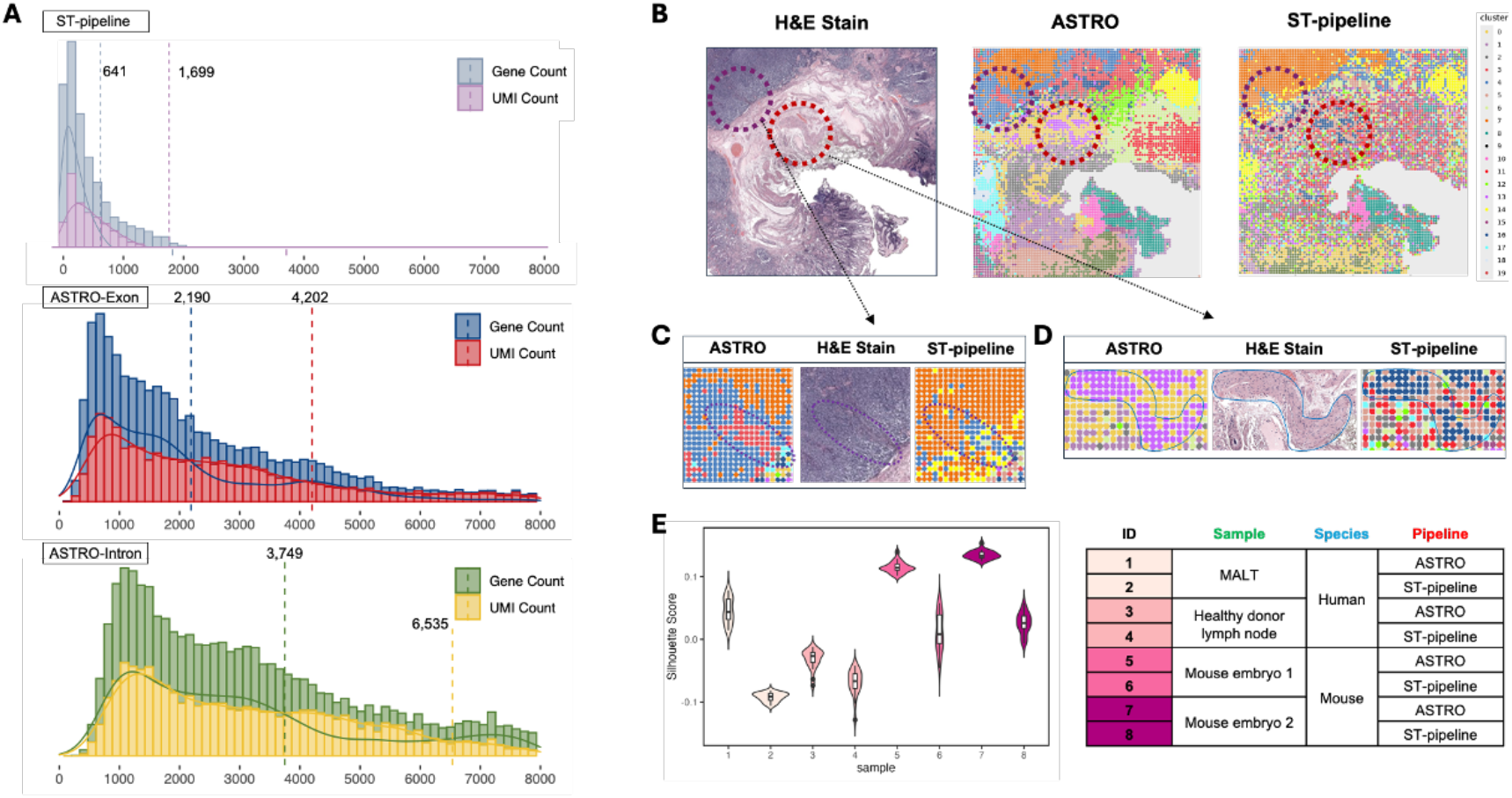
Benchmarking of downstream analysis based on different pipelines. **(A)** Distribution of detected gene/UMI counts per spatial pixel from different sources. Dashed lines indicate the average levels of gene or UMI counts. **(B)** Comparison of H&E images, ASTRO-based clustering, and ST-pipeline-based clustering for the MALT sample. The purple and red circles highlight two subtle structures in the tissue. **(C)** Enlarged view of the region within the purple circle in (B). **(D)** Enlarged view of the region within the red circle in (B). **(E)** Quantitative measurement of downstream analyses. Four samples were analyzed separately using ASTRO and ST-pipeline. Silhouette scores for each condition are shown.

## 4 Conclusions and Discussion

In this study, we developed the ASTRO pipeline and demonstrated its utility and performance across multiple datasets addressing diverse biological questions. Overall, ASTRO effectively quantifies on the spatial level the whole transcriptome, including non-coding RNAs, while capturing both RNA isoform details and RNA maturation stages in a spatial context. To the best of our knowledge, it is the first tool specifically designed for this purpose. Unlike most previous studies, which analyzed non-coding RNAs separately using different references and mapping steps, our pipeline examines all non-coding RNAs simultaneously, thereby streamlining downstream analyses.

In addition, our pipeline is specialized for FFPE samples, making it particularly useful for spatial profiling using clinical archives. This enhancement, together with broader RNA species coverage, improves downstream analyses when using FFPE datasets. This is especially important in clinical settings, as many FFPE samples have been stored under suboptimal conditions for years(Matsunaga *et al*., 2022). Consequently, maximizing the amount of sequenced information is critical for reliable analysis of FFPE sample sequencing.

Although this pipeline focuses on FFPE spatial transcriptome data, ASTRO is also compatible with non-FFPE spatial transcriptome datasets and single-cell sequencing data. Furthermore, if these datasets provide whole-transcriptome coverage (e.g., VASA-seq or STRS) (McKellar *et al*., 2023; Salmen *et al*., 2022), ASTRO can achieve the quantification of various RNA species in the entire transcriptome.

## Author Contributions

D.Z. and Z.B. designed the study. D.Z. Z.B. and Z.C. performed benchmarking analyses. D.Z. and Y.H. wrote the pipeline scripts, D.Z., Y.H and Z.C. maintained the git hub. M.G., J.L., and R.F. supervised the study. D.Z., Z.C, Z.B., J.L., and M.G. wrote the manuscript.

