## Supplementary figures and text for "ASTRO: Automated Spatial Whole-Transcriptome RNA-Expression Workflow"

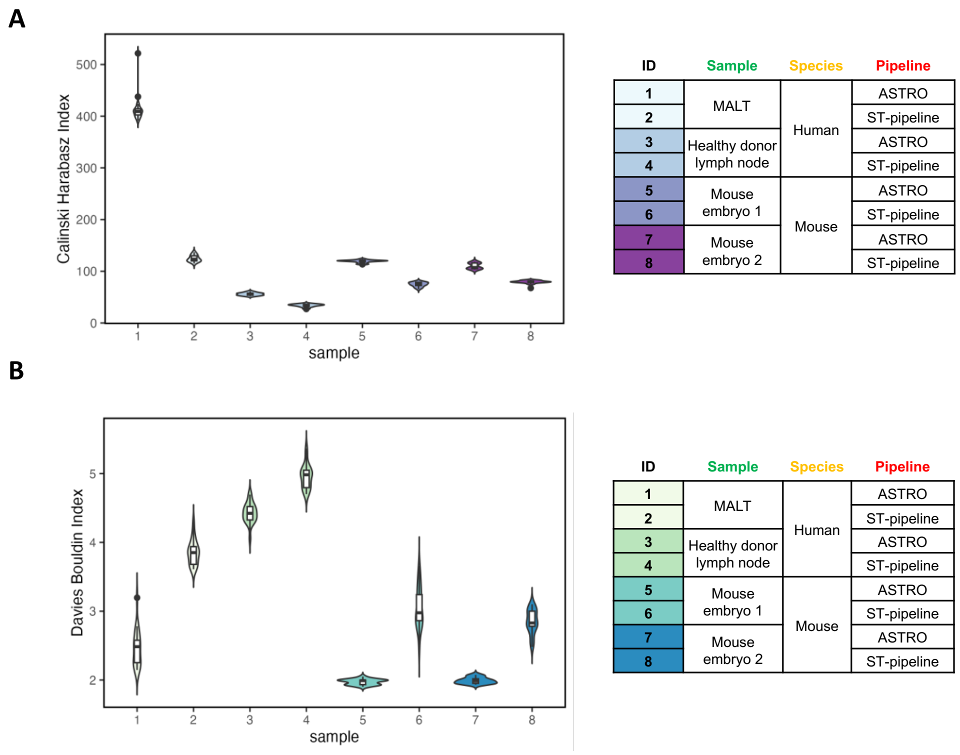


**Fig. S1** Quantitative measurement of downstream analysis. Four samples are analyzed by ASTRO and ST-pipeline, separately. The performance is shown as Calinski Harabasz Index **(A)** Davies Bouldin Index **(B)**, separately.

### *Data Subsampling and Benchmarking of Pipelines*

The original expression data matrices were randomly downsampled to 50% of the total reads, with subsampling repeated 20 times for each sample. Downstream gene expression analysis was conducted using the Seurat V5 pipeline (Hao et al., 2023). SCTransform (Choudhary & Satija, 2022; Hafemeister & Satija, 2019) was employed to normalize gene expression for each pixel, followed by principal component analysis (PCA) via the RunPCA function, retaining the top 50 principal components for further analysis. A K-nearest neighbor graph was then constructed using the FindNeighbors function, based on Euclidean distance in the PCA space, and pixel clustering was performed with the FindClusters function. To facilitate comparisons, custom script was deployed to fix the number of clusters at 15 in each round for both ASTRO-derived and ST-pipeline-derived data. Clustering performance was assessed using the Silhouette score, Calinski–Harabasz index, and Davies–Bouldin index, with the functions from scikit-learn. These metrics, which quantify intra-cluster cohesion and inter-cluster separation, are widely used to evaluate the quality of single-cell expression data clustering (Buitinck et al., 2013; Jiang et al., 2018; Leng et al., 2022; Møller & Madsen, 2023; Yu et al., 2022).

***Validation of Genomic Features***

ASTRO first identifies all reads mapped to a given feature. The pipeline then compares the read coverage in these regions to that of an extended background. Specifically, it counts the number of read start and end sites within an extended feature region (by default, 5 bp upstream and downstream, Region 1) and compares these counts to those in a larger background region (an additional 5 bp upstream and downstream, Region 2). A feature is considered valid if its coverage is significantly enriched relative to the background. The statistical method employed is a two-sample Poisson intensity rate comparison (test_poisson_2indep in the statsmodels package) (Gu et al., 2008; Ng et al., 2007; Seabold & Perktold, 2010). The test compares the densities of 5' and 3' read ends in Regions 1 and 2. When performing test_poisson_2indep, the number of 5'/3' reads located in Region 1 is assigned to count1, and the length of Region 1 is assigned to exposure1. Likewise, the number of start/end sites in Region 2 is assigned to count2, and the length of Region 2 is assigned to exposure2. A read is counted in Region 1 only if both its 5' and 3' ends fall within Region 1. A feature is considered invalid if either the 5' or 3' read ends within Regions 1 and 2 fail to satisfy the following criteria: (1) the density in Region 1 is at least $x$-fold higher than in Region 2, and (2) the $p$-value is below $y$. By default, $x = 2$ and $y = 0.05$.

***Spatial Presentation of the MALT Sample:***

The analysis was conducted using the Seurat V5 pipeline (Hao et al., 2023). SCTransform (Choudhary & Satija, 2022; Hafemeister & Satija, 2019) was employed to normalize gene expression for each pixel, followed by PCA via the RunPCA function. Elbow plots were used to determine the number of principal components to retain: 10 for ASTRO-based data and 6 for ST-pipeline-based data. A K-nearest neighbor graph was then constructed using the FindNeighbors function, based on Euclidean distance in the PCA space, and pixel clustering was performed with the FindClusters function. To facilitate comparisons, the resolution of clusters was set to 1.2 for both ASTRO- and ST-pipeline-derived data. Clustering performance was assessed using the Silhouette score, Calinski–Harabasz index, and Davies–Bouldin index, with functions from scikit-learn (Buitinck et al., 2013; Jiang et al., 2018; Leng et al., 2022; Møller & Madsen, 2023; Yu et al., 2022). These metrics, which quantify intra-cluster cohesion and inter-cluster separation, are widely used to evaluate the quality of single-cell expression data clustering.

***References:***

Buitinck, L., Louppe, G., Blondel, M., Pedregosa, F., Müller, A. C., Grisel, O., Niculae, V., Prettenhofer, P., Gramfort, A., Grobler, J., Layton, R., Vanderplas, J., Joly, A., Holt, B., & Varoquaux, G. (2013). API design for machine learning software: experiences from the scikit-learn project. https://arxiv.org/abs/1309.0238v1

Choudhary, S., & Satija, R. (2022). Comparison and evaluation of statistical error models for scRNA-seq. Genome Biology, 23(1), 1–20. https://doi.org/10.1186/S13059-021-02584-9/FIGURES/4

Gu, K., Ng, H. K. T., Man, L. T., & Schucany, W. R. (2008). Testing the ratio of two poisson rates. Biometrical Journal. Biometrische Zeitschrift, 50(2), 283–298. https://doi.org/10.1002/BIMJ.200710403

Hafemeister, C., & Satija, R. (2019). Normalization and variance stabilization of single-cell RNA-seq data using regularized negative binomial regression. Genome Biology, 20(1), 1–15. https://doi.org/10.1186/S13059-019-1874-1/FIGURES/6

Hao, Y., Stuart, T., Kowalski, M. H., Choudhary, S., Hoffman, P., Hartman, A., Srivastava, A., Molla, G., Madad, S., Fernandez-Granda, C., & Satija, R. (2023). Dictionary learning for integrative, multimodal and scalable single-cell analysis. Nature Biotechnology 2023 42:2, 42(2), 293–304. https://doi.org/10.1038/s41587-023-01767-y

Jiang, H., Sohn, L. L., Huang, H., & Chen, L. (2018). Single cell clustering based on cell-pair differentiability correlation and variance analysis. Bioinformatics (Oxford, England), 34(21), 3684–3694. https://doi.org/10.1093/BIOINFORMATICS/BTY390

Leng, D., Zheng, L., Wen, Y., Zhang, Y., Wu, L., Wang, J., Wang, M., Zhang, Z., He, S., & Bo, X. (2022). A benchmark study of deep learning-based multi-omics data fusion methods for cancer. Genome Biology, 23(1). https://doi.org/10.1186/S13059-022-02739-2

Møller, A. F., & Madsen, J. G. S. (2023). JOINTLY: interpretable joint clustering of single-cell transcriptomes. Nature Communications 2023 14:1, 14(1), 1–15. https://doi.org/10.1038/s41467-023-44279-8

Ng, H. K. T., Gu, K., & Tang, M. L. (2007). A comparative study of tests for the difference of two Poisson means. Computational Statistics & Data Analysis, 51(6), 3085–3099. https://doi.org/10.1016/J.CSDA.2006.02.004

Seabold, S., & Perktold, J. (2010). statsmodels: Econometric and statistical modeling with python. 9th Python in Science Conference.

Yu, L., Cao, Y., Yang, J. Y. H., & Yang, P. (2022). Benchmarking clustering algorithms on estimating the number of cell types from single-cell RNA-sequencing data. Genome Biology, 23(1), 1–21. https://doi.org/10.1186/S13059-022-02622-0/TABLES/1
